# Chromosome-level genome assembly and transcriptome- based annotation of the oleaginous yeast *Rhodotorula toruloides* CBS 14

**DOI:** 10.1101/2021.04.09.439123

**Authors:** Giselle C. Martín-Hernández, Bettina Müller, Mikołaj Chmielarz, Christian Brandt, Martin Hölzer, Adrian Viehweger, Volkmar Passoth

## Abstract

*Rhodotorula toruloides* is an oleaginous yeast with high biotechnological potential. In order to understand the molecular physiology of lipid synthesis in *R. toruloides* and to advance metabolic engineering, a high-resolution genome is required. We constructed a genome draft of *R. toruloides* CBS 14, using a hybrid assembly approach, consisting of short and long reads generated by Illumina and Nanopore sequencing, respectively. The genome draft consists of 23 contigs and 3 scaffolds, with a N50 length of 1,529,952 bp, thus largely representing chromosomal organization. The total size is 20,534,857 bp with a GC content of 61.83%. Transcriptomic data from different growth conditions was used to aid species-specific gene annotation. In total we annotated 9,464 genes and identified 11,691 transcripts. Furthermore, we demonstrated the presence of a potential plasmid, an extrachromosomal circular structure of about 11 kb with a copy number about three times as high as the other chromosomes.

**Significance:** This obtained high-quality draft genome provides the suitable framework needed for genetic manipulations, and future studies of lipid metabolism and evolution of oleaginous yeasts. The identified extrachromosomal circular DNA may be useful for developing efficient episomal vectors for the manipulation of *Rhodotorula* yeasts.

**Data deposition:** This project has been deposited at ENA under the accession PRJEB-40807.

## Introduction

The basidiomycete yeast *Rhodotorula toruloides* is an oleaginous microorganism with high biotechnological potential for lipid and carotenoid production. This yeast can naturally accumulate lipids up to 70% of dry cell weight, and a number of carotenoids (González-García et al., 2017; Park et al., 2018). *R. toruloides* can be cultivated to high cell densities on a wide range of substrates, including lignocellulose hydrolysate and other residual products (Júnnior et al., 2020; Sànchez Nogué et al., 2018; Singh et al., 2018; Wiebe et al., 2012, Chmielarz et al. 2021). This makes *R. toruloides* a promising host for the production of single-cell oil, as a sustainable and less controversial alternative to plant-derived oils for the production of biofuels, food and feed additives (Park et al., 2018).

The molecular physiology behind lipid synthesis in *R. toruloides* has been relatively little explored, which hinders effective metabolic engineering for improved lipid production. Some draft genome sequences from *R. toruloides* strains have been determined using short-read sequencing technologies, however a complete picture of the *R. toruloides* genome on chromosomal level is still lacking (Hu & Ji, 2016; Kumar et al., 2012; Morin et al., 2014; Paul et al., 2014; Sambles et al., 2017; Tran et al., 2019; Zhang et al., 2016; Zhu et al., 2012). The combination of short- and long-read sequencing strategies has been shown to improve the accuracy of genome sequences in yeast (Olsen et al., 2015; Tiukova et al., 2019).

Thus, we combined Nanopore long-read sequencing and Illumina short-read sequencing to obtain a chromosome-level genome assembly. We further generated comprehensive transcriptomic data from different growth conditions to aid species-specific annotation. The results provide a valuable resource for pathway analysis and manipulation of *R. toruloides* and enable better understanding of genome biology and evolution of basidiomycetous yeasts.

## Materials and Methods

### DNA extraction

*R. toruloides* CBS 14 (Westerdijk Fungal Biodiversity Institute, Utrecht, the Netherlands) was grown in 50 ml YPD medium (Chmielarz et al., 2019). Cells were harvested during exponential growth and the cell wall was digested according to Pi et al. (2018). DNA was extracted from the protoplasts using NucleoBond® CB 20 Kit (Macherey-Nagel, Germany). Concentration, purity and integrity were assessed with Qubit™ 4 Fluorometer, NanoDrop® ND-1000 Spectrophotometer (Thermo Fisher Scientific, USA) and agarose gel electrophoresis, respectively.

### RNA extraction

*R. toruloides* was cultivated in 500 mL bioreactors (Multifors, Infors HT, Bottmingen, Switzerland) with either a mixture of lignocellulose hydrolysate and crude glycerol or crude glycerol only as carbon source (Blomqvist et al. 2018; Chmielarz et al., 2021). Additionally, the media contained 0.75 g/L yeast extract (BactoTM Yeast Extract, BD, France), 1 g/L MgCl_2_ 6xH_2_O (Merck KGaA, Germany), 2 g/L (NH_4_)_2_HPO_4_ (≥98%, Sigma-Aldrich, USA), and 1.7 g/L YNB without amino acids and ammonium sulfate (DifcoTM, Becton Dickinson, France). Cultivations were performed in triplicates, at 25°C, pH 6 and oxygen tension of 21%.

Total RNA was extracted in triplicates from 5 mL samples withdrawn after 10, 40, 72 and 96 h, respectively from bioreactors with mixed carbon source, and 10, 30 and 60 h, respectively from bioreactors with glycerol as sole carbon source, using the Quick-RNA(tm) Fecal/Soil Microbe MicroPrep Kit (Zymo Research, USA) according to the manufacturer’s instructions with some modifications (Manzoor et al., 2016). Briefly, the cells were harvested and resuspended in 1 mL Trizol and disrupted in a FastPrep -24 bead beater (M.P. Biomedicals, USA) at speed 6.0 m/s for 40s. The homogenate was separated into layers by adding 0.2 mL chloroform and centrifugation. 400 µl of the upper layer containing the RNA was further processed as described in the manufacturer’
ss manual. DNAse I treatment was performed as described in the manual applying 26 U/mL at 37° for 15 minutes. The technical replicates of the retrieved RNA were pooled prior rRNA depletion, which was performed using the human riboPOOL™ (siTOOLsBiotech, Germany) and Streptavidin-coated magnetic beads (Thermo Fisher Scientific, Norway) following the two-step protocol of the manufacturer. rRNA-depleted samples were purified by ethanol precipitation. RNAse inhibitor (1 U/µL, Thermo Fisher) was added before storage. Total RNA and rRNA-depleted samples were tested for integrity and quantity on RNA Nano Chips (Agilent 2100 Bioanalyzer System, Agilent Technologies, Germany).

### Library preparation and sequencing

The extracted DNA was divided into two samples and sequenced using either MinION (Oxford Nanopore Technologies) or Illumina sequencing platform. Before Nanopore library preparation, 5 µg of the retrieved DNA were “pre-cleaned” using 31.5 µL of AMPure magnetic beads, for removing short DNA fragments (Brandt et al., 2019). The DNA library was prepared using the Ligation Sequencing Kit SQK-LSK109 (Oxford Nanopore Technologies, Oxford, UK) and a modified protocol (Brandt et al., 2019). The library was loaded onto a FLO-MIN106 flow cell attached to the MinION device (Oxford Nanopore Technologies). Sequencing was performed using the MinKNOW software (Oxford Nanopore Technologies) according to the protocol from Brandt et al. (2019).

Short-read paired-end sequencing of DNA and rRNA-depleted samples was performed on the Illumina Novaseq platform (S prime, 2x 150 bp) using TruSeq PCR free DNA library preparation kit (Illumina Inc.).

In order to verify the circular structure of contig_63, Sanger sequencing (Macrogen Europe B.V, the Netherlands) was performed from PCR amplicons using the primers shown in supplementary table s1. The PCR mixture consisted of 0.3 µL DNA, 1.25 µL primer, 12.5 µL Dream Taq Green PCR Mix (Thermo Scientific, Lithuania) and 0.8 µL DMSO in a total volume of 25 µL. Amplification was conducted as follows: initial denaturation at 95°C for 5 min, 35 cycles of denaturation (95°C for 30 s), annealing (1 min) and elongation (72°C) followed by a final extension at 72°C for 10 min. Annealing temperatures and elongation times were adapted to the respective primer combination (table s2). Amplification was assessed on agarose gel electrophoresis at 8.2 V/cm electric field strength. The gel was prepared using GelRed® Nucleic Acid Gel Stain (Biotium), 1.0% agarose and TAE 1X buffer (VWR Life Science). The PCR products from the corresponding sizes were purified using GeneJet Gel Extraction kit (Thermo Scientific, Lithuania). Geneious prime version 2021.0.1 (Biomatters Ltd.) was used for assembly of the Sanger sequences obtained and to align *FAS2* and *FAS21* from *R. toruloides* CBS 14 and *FAS2* from *R. toruloides* NP11.

### Quality control and genome assembly

Short and long reads generated by Illumina and Nanopore sequencing, respectively, were combined for hybrid *de-novo* assembly. Before, the quality of the long reads was ensured using NanoPlot v1.25.0 (De Coster et al., 2018) and all short reads were removed until reaching a target base coverage with Filtlong v0.2.0 (https://github.com/rrwick/Filtlong) (Wick, n.d.) applying the parameters --target_bases 5000000000 and --length_weight 8. Short reads were quality-trimmed and adapter-clipped with fastp v0.20.0 (Chen et al., 2018) using parameters - 5 -3 -W 4 -M 20 -l 25 -x -n 5 -z 6. To achieve high contiguity in the initial assembly step, a draft assembly was produced from the preprocessed long reads using Flye v2.8 (Kolmogorov et al., 2019), setting the suggested genome size to 20 Mbp and keeping the plasmid and meta options activated. The Flye draft assembly was further subjected to long-read-based polishing using two rounds of Racon v1.4.7 (Vaser et al., 2017) followed by one round of Medaka v0.10.0 (https://github.com/nanoporetech/medaka) (Technologies, n.d.) using the model r941_min_high. We used minimap2 (-x map-ont) v2.17 (Li, 2018) and samtools v.1.10 (Li et al., 2009) to prepare alignment files for the Racon polishing. The assembly graph was visualized and investigated via Bandage v0.8.1 (Wick et al., 2015). The long-read assembly was finally subjected to two Pilon v1.23 (Walker et al., 2014) rounds, in which the quality-filtered short reads were used for final polishing. Alignment files for Pilon were prepared using BWA v0.7.17 (Li & Durbin, 2009) and samtools. We mapped the Illumina and Nanopore reads with BWA and minimap2, respectively, to the final assembly and used the pileup.sh script from BBMap v38.86 (Bushnell, 2014) to calculate coverage histograms for each contig with a bin size of 1000 nt and plotted them with ggplot2. Besides the visual inspection of coverage patterns, we used the samtools coverage function and the faidx command on the long-read-mapped data to filter out contigs violating at least one of the following cutoffs: minimum read number = 100, minimum bases covered by reads = 5000, minimum read coverage = 15, and minimum read coverage depth (amount of bases covered by reads in comparison to contig length) = 10. The resulting quality-controlled and cleaned assembly file was used for annotation and chromosome analyses.

### Annotation

The final assembly was first screened for repetitive regions using RepeatMasker v4.0.9 (Nishimura, 2000), checked for completeness using BUSCO v3.0.2 (Simão et al., 2015; Waterhouse et al., 2018) with the fungi_odb9 database as reference and visualized using chromoMap v0.2 (Anand, 2019). Contigs, larger than 10 kb but smaller than 250 kb were checked for circularization using a python script (https://github.com/Kzra/Simple-Circularise, v1.0) (Kitson, n.d.).

Taxonomic classification was performed using sourmash v2.0.1 (Pierce et al., 2019) and its “LCA” method. Both indexing the tree and querying genomes used a k-mer size of 31 and a sampling fraction of 10-4. The LCA index was derived from publicly available genomes (GenBank, https://osf.io/4f8n3/).

The repeat-masked assembly was further annotated using a combination of homology-based gene comparison and RNA-Seq-derived transcript information. First, we used MetaEuk v1.ea903e5 (Karin et al., 2020) with the “easy-predict” subcommand for draft gene annotation providing all proteins (filtered models; best) obtained from the JGI fungal genome portal MycoCosm (http://jgi.doe.gov/fungi; downloaded April 2020) as a database. Next, we combined and improved the MetaEuk annotation with the obtained RNA-Seq data. Prior to this, RNA-Seq datasets were quality-checked using FastQC v0.11.8 (https://www.bioinformatics.babraham.ac.uk/projects/fastqc/) (Andrews et al., 2010) and trimmed and adapter-clipped using fastp with the aforementioned parameters. Potential residual rRNA was removed by SortMeRNA v2.1b (Kopylova et al., 2012). The processed RNA-Seq reads were per-sample-mapped to the assembly using HISAT v2.2.0 (Kim et al., 2015) and subsequently provided to StringTie v2.1.1 (Kovaka et al., 2019; Pertea et al., 2015) for transcript annotation, guided by the initial MetaEuk gene annotation. Finally, the MetaEuk-guided StringTie information from each individual RNA-Seq sample were merged into a final single annotation using the StringTie merge functionality. In this step, we again provided the full MycoCosm-based MetaEuk annotation next to the individual RNA-Seq annotations to also include genes that were not recovered in the RNA-Seq data. Lastly, we extracted all annotated gene sequences and used the dammit pipeline v1.2 (http://www.camillescott.org/dammit) (Scott, n.d.) for functional annotation. Dammit unites different databases for annotation: Pfam-A, Rfam, OrthoDB, and again BUSCO (fungi_odb9) that we have decorated with additional information derived from UniProt/Swiss-Prot. We extracted the GO terms annotated by dammit and counted their appearance to apply a weight per GO term. GO terms that occured at least 10 times were subjected to REVIGO (https://journals.plos.org/plosone/article?id=10.1371/journal.pone.0021800) to summarize them by removing redundancy and to visualize the results in semantic similarity-based scatterplots and interactive graphs.

We used a custom python script to identify potential telomeres by using known motifs at contig ends. This information was provided along with the Nanopore reads to Tapestry v1.0.0 (Davey et al., 2020).

## Results and Discussion

The hybrid *de-novo* assembly for *R. toruloides* CBS 14 (fig. 1) resulted in a 20,534,857 bp genome, which is in line with the 20.4 Mb median reported for this and other *R. toruloides* strains (table 1). The overall GC content of the obtained genome is 61.83%, which corresponds to previously reported values (61.9% on average). The identified repetitive sequences represent 2.01% of the total genome length, of which 1.56% are single repeats and 0.45% low complexity regions. The assembled genome resulted in 23 contigs and 3 scaffolds ranging from 5,778 to 1,965,970 bp, and an N50 length of 1,529,952 bp (fig. 2). This number of contigs and scaffold is significantly lower than that obtained using short-read sequencing only: the lowest number achieved in previous studies is 186 contigs (table 1). This clearly shows the higher accuracy, contiguity and completeness of the genome presented here, achieved through the improved coverage using a hybrid assembly approach. 56.37% of the genome is represented by 4 contigs and 3 scaffolds, each larger than 1 Mb. The rest is allocated in 10 medium-size contigs (0.5 - 1 Mb) and in 9 small contigs (< 0.5 Mb), four of which are circularized. Telomere sequences could be detected at both termini of two contigs and at one terminus each of 15 contigs (fig. 2). De Jonge et al. (1986) identified at least 11 chromosomes in *R. toruloides* CBS 14 by pulsed field gel electrophoresis (PFGE). Zhu et al. (2012) predicted 16 chromosomes in *R. toruloides* NP11 applying Illumina sequencing and gap closing using genome walking. When comparing the size of the obtained contigs and scaffolds with the corresponding bands found by PFGE, we identified at least 18 chromosomes in *R. toruloides* CBS 14.

**Table 1.**
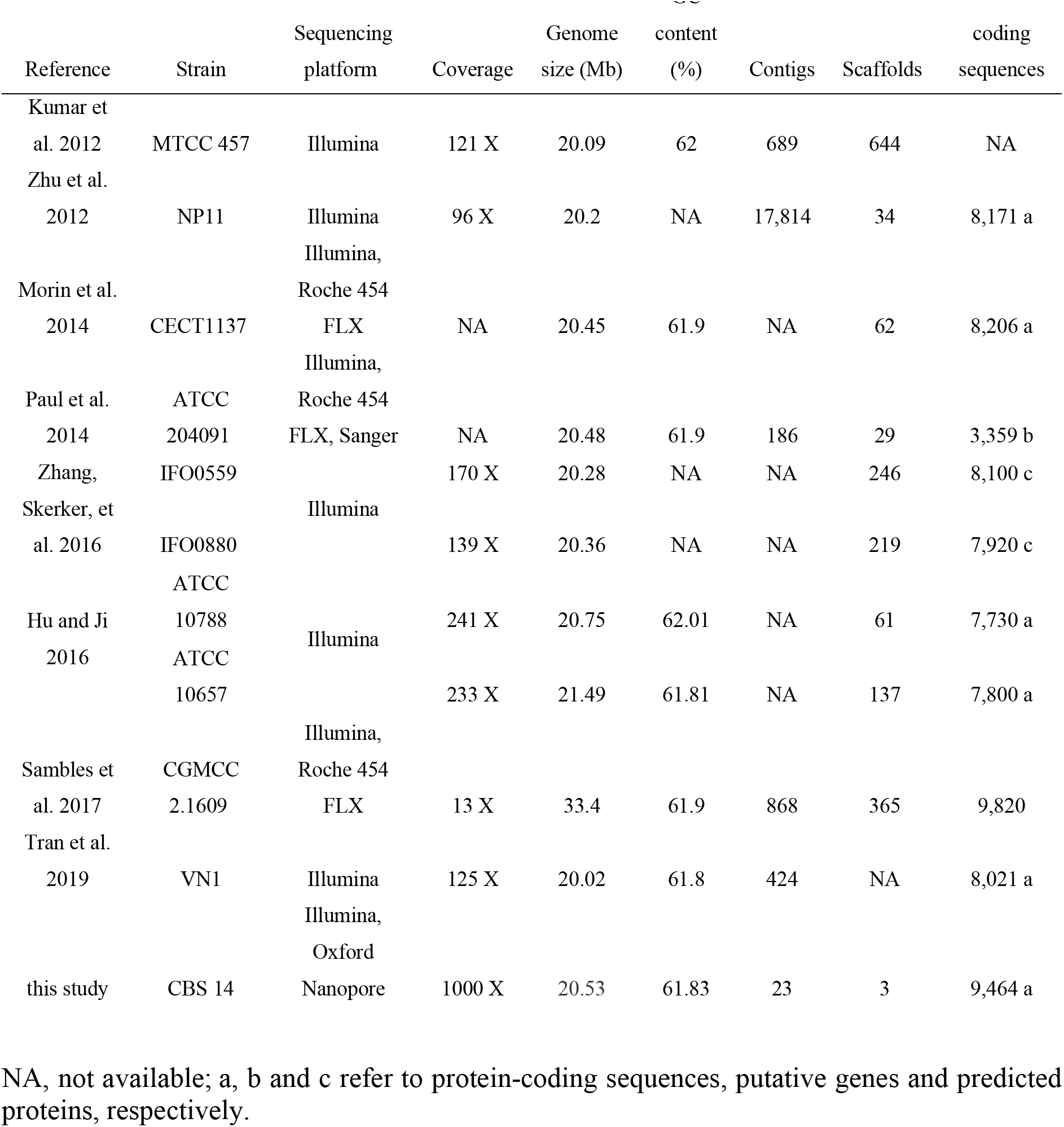
Genome assembly statistics of *Rhodotorula toruloides* strains and sequencing platforms used.

**Fig. 1.**
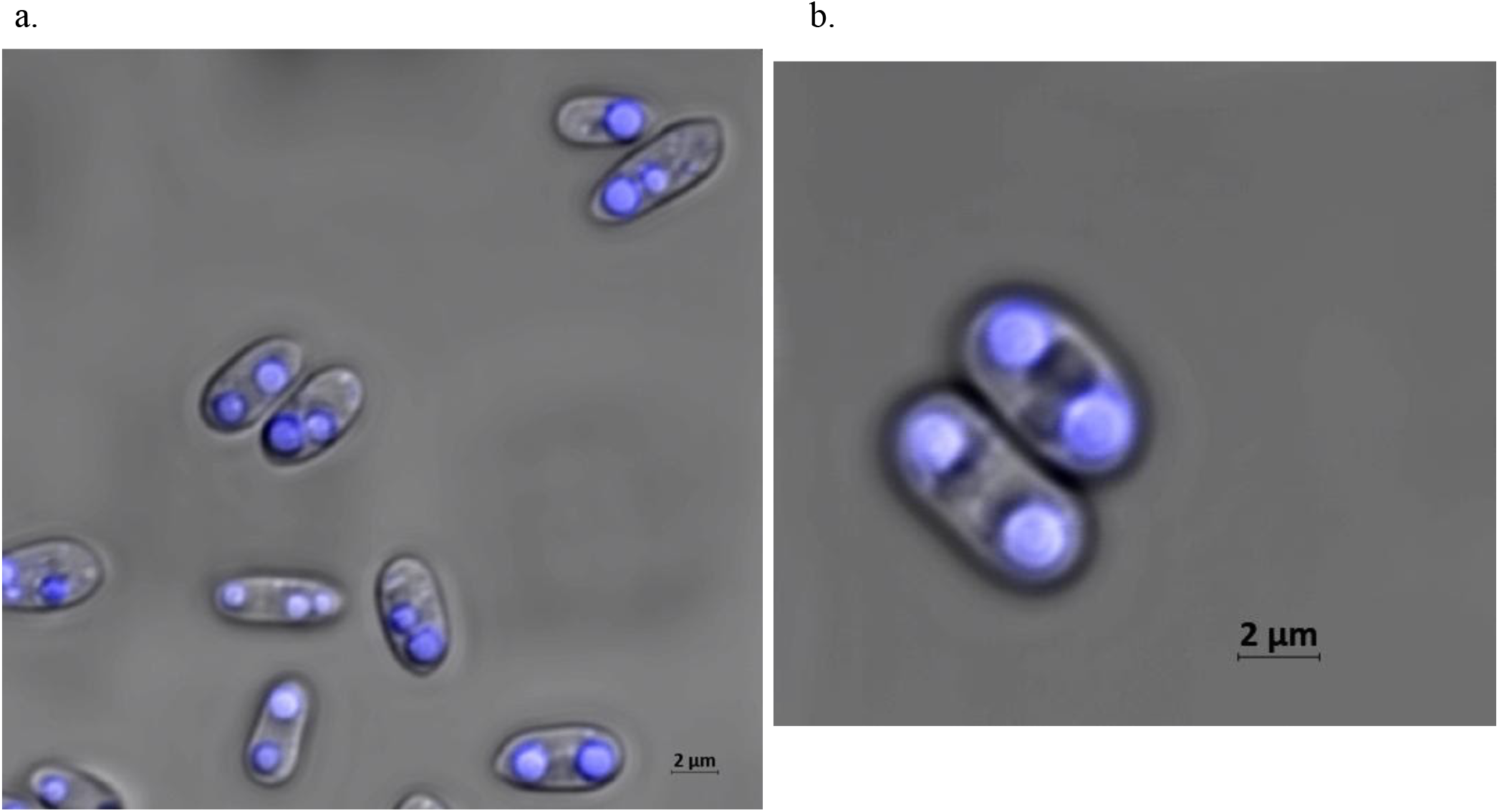
Confocal microscope picture of Nile Red-stained lipid droplets of *Rhodotorula toruloides* CBS 14. (a) Group of cells at stationary phase (b) Magnified view of cells.

**Fig. 2.**
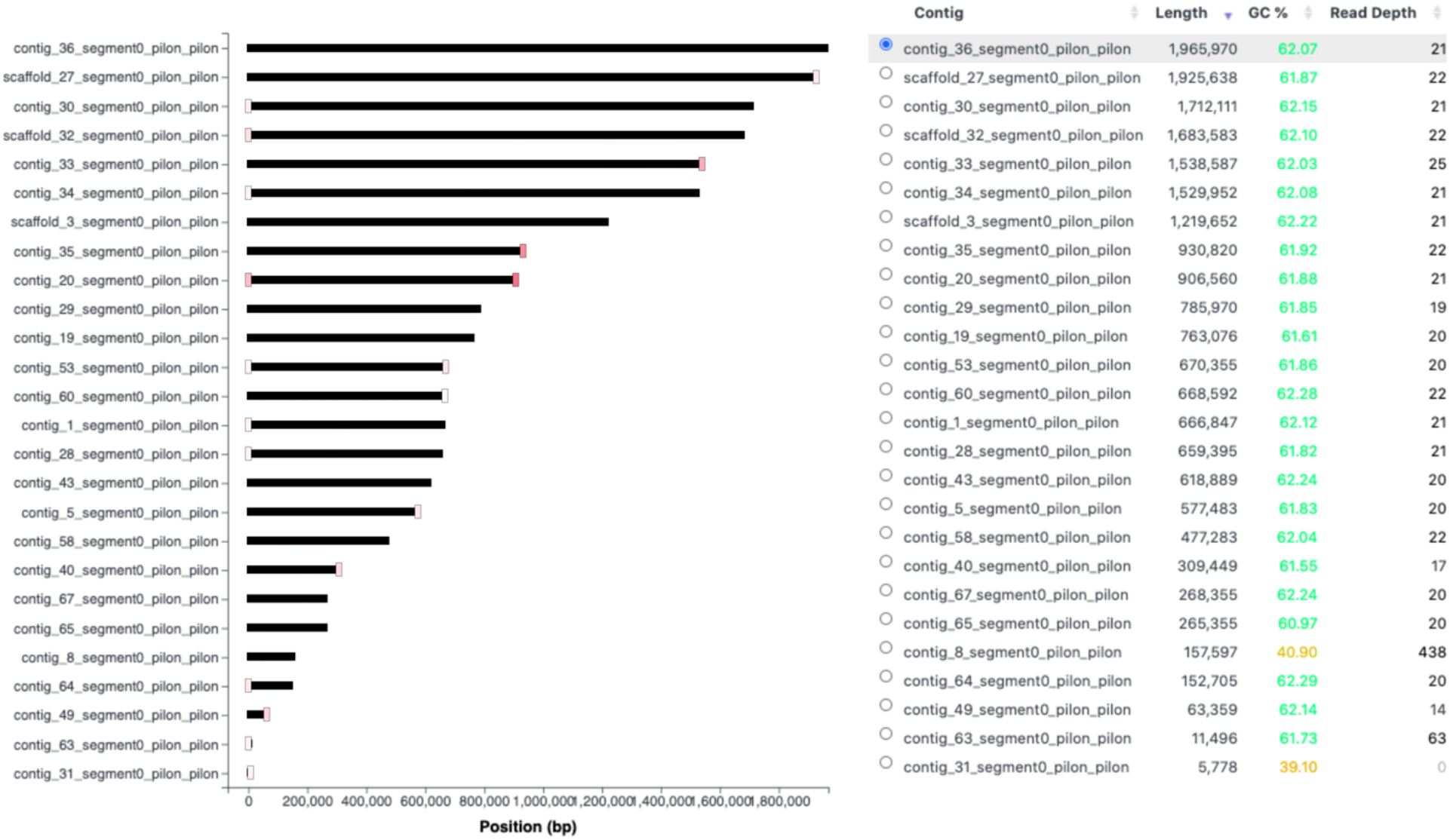
Overview of the genome assembly of *Rhodotorula toruloides* CBS 14 including contig and scaffold lengths, GC contents and obtained read depths. Telomere sequences are represented with vertical lines at the end of contigs. Line transparency from red to gray indicates their number in decreasing order.

The mitochondrial genome was recovered in one contig with a length of 157 kb. However, the final step of circularizing sequences in an assembly remains challenging and thus can result in shorter or even longer contigs, since assembly programs cannot clearly define where a circular DNA ends, even despite longer reads (Hunt et al. 2015). Thus, tandem repeats at the end of a contig can increase the length of a linear representation of a circular sequence. Such repeats can be identified by overlapping sequences at the ends of a contig. To account for this, in an additional step, we aligned the contig against itself to identify and remove circular repeats, ultimately resulting in an actual mitochondrial genome size of 69 kb. The mitochondrial genome has a GC content of 40.9%, which agrees with the GC values previously reported for *R. toruloides* strains NP11 and NBRC 0880 (Zhou et al. 2020; Coradetti et al. 2018). In addition to the mitochondrial genome, three further circular contigs (contig_64, contig_49, contig_63) were predicted. Among these, Contig 63 was particularly noticeable because it showed a read depth approximately three times higher than the other chromosomes, which may indicate relaxed replication regulation (fig. 2). We confirmed the circular structure of Contig 63 by overlapping amplification using sequence-specific primers and subsequent Sanger sequencing of the amplicons obtained (supplementary fig. S1, supplementary file 2). This showed that the circular structure is about 1,143 bp larger than the one predicted (supplementary fig. S1). As explained above, circular sequence structures possess a certain challenge for assembly tools and can particularly result in longer but also shorter contigs that miss parts of the actual sequence at the end of a contig (Hunt et al. 2015). Extrachromosomal endogenous DNA have been previously found in *Saccharomyces cerevisiae* (Strope et al. 2015; Møller et al. 2015). The presence of DNA mitochondrial plasmids in filamentous fungi, including some Basidiomycota species, have also been widely acknowledged before (Griffiths 1995; Wang et al. 2008; Cahan & Kennell 2005). Within them, mitochondrial circular DNA plasmids are less frequently found than linear. They are typically within a length range of 2.5–5 kb, and encoding enzymes involved in their replication such as a DNA polymerase or a reverse transcriptase (Cahan & Kennell 2005; Wang et al. 2008). The genes annotated in Contig 63 are *UTP22, H2A* and *H2B* which are encoding for RNA-associated protein 22, Histone H2A and Histone H2B, respectively (supplementary table s3). A truncated copy of *UTP22* from *R. toruloides* NP11 was also annotated in Contig 63 and identified through Blast search (supplementary table s3). The CDS of *UTP22* and *H2A* are also found in another contig within the genome. Both of the *UTP22* genes encoded in contig 63 are shortened compared to the *UTP22* gene that we also found encoded on scaffold 32. The *UTP22* gene on scaffold 32 shares 100% sequence identity with the *UTP22* gene identified in strain NP11. Furthermore, *S. cerevisiae* ACS-like sequences described by Dhar et al. (2012) are not present in Contig 63. AT-rich ACS elements of ARS have been found to be degenerate and in *Schizosaccharymces pombe*, which is closer related to *S. cerevisiae* than *R. toruloides*, the absence of such consensus sequence has been reported (Dhar et al. 2012; Iwakiri et al. 2005). This is the first time that such extrachromosomal circular DNA has been detected in Basidiomycetes. Such structures may be useful for developing efficient episomal vectors for the manipulation of *Rhodotorula* yeasts.

Analysis of the RNA-Seq data of *R. toruloides* CBS 14 using StringTie software and guided by the MetaEuk annotation (based on all protein sequences from MycoCosm) resulted in the annotation of 9,464 genes with 11,691 transcripts. This number is significantly higher than previously reported for *R. toruloides* genomes (table 1). Five transcripts were annotated within the circular Contig 63 (supplementary table s3). The higher number of transcripts than protein-coding genes can be explained by alternative splicing and non-coding RNAs. Zhu et al. (2012) found 1,371 genes encoding two or more transcript isoforms in *R. toruloides* NP11. The average number of exons per gene is 5.95 in *R. toruloides* CBS 14 (supplementary table s4). Confirming the findings of Zhu et al. (2012), we observed a predominance of split genes in the genome, with a total of 8,550 for *R. toruloides* CBS 14. After functional annotation of coding sequences through Gene Ontology (GO) terms we clustered annotations into the categories: biological processes, cellular components and molecular functions (supplementary fig. S2-S4). Genes encoding enzymes crucial for lipid and carotenoid metabolisms such as *CDC19, MAE1, MAE2, ACL1, ACC1, FAS1, FAS2, OLE1, ACAD10, ACAD11, crtYB, crtI* and *BTS1* are present in the genome (supplementary table s5). Similar to the situation in *S. cerevisiae* (Burkl et al., 1972), the genes encoding the *α*- and β-subunits of FAS are located in different chromosomes (supplementary table s5).

The MetaEuk annotation identified a gene (*FAS21*) on the opposite strand of the *FAS2*-gene (supplementary fig. S5), which encodes the *α*-subunit of the fatty acid synthase complex (FAS) in *R. toruloides* NP11 (Zhu et al., 2012). *FAS21* contains 2 exons and its mRNA would be complementary to parts of the *FAS2*-sequence. There is a growing number of natural antisense transcripts identified in fungal transcriptome analyses (Chacko & Lin, 2013; Donaldson & Saville 2012). *FAS21* transcript might be involved in controlling *FAS2* expression. However, we did not identify a *FAS21* transcript under the experimental conditions carried out, thus the involvement of *FAS21* in the regulation of *FAS2*-expression and thus fatty acid synthesis still needs to be verified. Three different transcripts of *FAS2* were identified containing 3, 12 and 16 exons, respectively (supplementary table s6).

BUSCO orthologs analysis showed that 96.9% of the assessed genes were identified and complete (96.6% single-copy and 0.3% duplicated), 1.7% fragmented and only 1.4% were lacking, demonstrating the high quality of the hybrid genome assembly (supplementary fig. S6).

In the case of the transcriptomic data, 84.1% of the genes were indicated as complete (69.3% single copy,14.8% duplicated), 14.1% as fragmented and 1.8% genes as missing.

The genome annotation and assembly correspond to the following taxonomic classification: Eukaryota superkingdom, Basidiomycota phylum, Microbotryomycetes class, Sporidiobolales order, Sporidiobolaceae family, *Rhodotorula* genus, *Rhodotorula toruloides* species, CBS 14 strain.

## Acknowledgements

The work was supported by the Swedish Research Council for Environment, Agricultural Sciences and Spatial Planning (Formas) [grant number 2018-01877]. Illumina sequencing was performed by the SNP&SEQ Technology Platform in Uppsala, which is part of the National Genomics Infrastructure (NGI) Sweden and Science for Life Laboratory. The SNP&SEQ Platform is also supported by the Swedish Research Council and the Knut and Alice Wallenberg Foundation. AV, CB and MH are shareholders of nanozoo GmbH.

## Authors’ contributions

Giselle C. Martín-Hernández: writing -original draft, investigation, formal analysis, visualization

Bettina Müller: writing -Review and editing, supervision, formal analysis, methodology Mikołaj Chmielarz: Investigation

Christian Brandt: resources, data curation, validation, writing-Review and editing,

Martin Hölzer: resources, data curation, validation, writing-Review and editing

Adrian Viehweger: resources, data curation, validation, writing-Review and editing

Volkmar Passoth: Conceptualization, writing -Review and editing, funding acquisition, project administration, resources

## Notes

### Competing Interest Statement

The authors have declared no competing interest.

