## Supplemental tables and figures for "Chromosome-level genome assembly and transcriptome- based annotation of the oleaginous yeast *Rhodotorula toruloides* CBS 14"

[1]Department of Molecular Sciences, Swedish University of Agricultural Sciences, Uppsala, Sweden, [2]nanozoo GmbH, Leipzig, Germany, [3]Institute for Infectious Diseases and Infection Control, Jena University Hospital, Jena, Germany, [4]RNA Bioinformatics and High-Throughput Analysis, Friedrich Schiller University Jena, Jena, Germany, [5]Department of Medical Microbiology, University Hospital Leipzig, Germany

Giselle C. Martín-Hernández and Bettina Müller equally contributed to this paper and shall both be regarded as first authors

**Supplementary Table 1.** Primers used for the validation of Contig 63 circular structure
**Supplementary Table 2.** PCR amplification conditions
**Supplementary Table 3.** Gene and transcript annotation in Contig 63
**Supplementary Table 4.** Distribution of exon counts in the transcriptome
**Supplementary Table 5.** Examples of lipid and carotenoid metabolism related genes in *Rhodotorula toruloides* CBS 14 genome assembly
**Supplementary Table 6.** Gene and transcript annotation of the $\alpha$- subunit of fatty acid synthase complex
**Supplementary Figure 1.** Validation of the circular structure of Contig 63.
**Supplementary Figure 2.** Gene Ontology (GO) term summaries belonging to the GO topic: molecular functions

**Supplementary Figure 3.** Gene Ontology (GO) term summaries belonging to the GO topic: biological processes

**Supplementary Figure 4.** Gene Ontology (GO) term summaries belonging to the GO topic: cellular components

**Supplementary Figure 5.** *De-novo* assembly of *FAS2* and *FAS21* of *Rhodotorula toruloides* CBS 14 with *FAS2* from *Rhodotorula toruloides* NP11

**Supplementary Figure 6.** A benchmark of near-universal single-copy orthologs assessed in the hybrid genome assembly

**Supplementary Table 1.** Primers used for validating the circular structure of Contig 63

| Primer | Sequences (5´- 3´) |
|---|---|
| 631F | CTCGGCCTCACTTCAGACTGGTG |
| 631R | CACCAGTCTGAAGTGAGGCCGAG |
| 633R | GTACACCGTTCTCAGACAGTGCGC |
| 635F | CTTCAACCCGTCCAATCCGCCTATC |
| 636R | CTCGCTCAACACGCTCGAGTTCAC |
| 638F | GCGACGACTTGTCCCTGTTCTTG |
| 638R | CAAGAACAGGGACAAGTCGTCGC |

**Supplementary Table 2.** Primer combinations and PCR conditions used for amplification of Contig63

| Fragment | Primer pairs | Annealing Temperature (°C) | Elongation time (min) |
|---|---|---|---|
| 1 | 631F-636R | 69 | 7 |
| 2 | 631F-633R | 69 | 2 |
| 3 | 635F-636R | 71 | 1 |
| 4 | 635F-638R | 67 | 3 |
| 5 | 638F-633R | 67 | 3 |
| 6 | 635F-631R | 69 | 3 |
| 7 | 638F-631R | 67 | 1 |
| 8 | 638F-636R | 67 | 7 |

**Supplementary Table 3.** Gene and transcript annotation of Contig 63

| | from | to | Sense | Gene ID | Transcript ID | Gene code | Name |
|---|---|---|---|---|---|---|---|
| gene | 13 | 429 | - | NANOZOOG6896 | | | |
| transcript | 13 | 429 | - | NANOZOOG6896 | NANOZOOT6896.1 | *UTP22* [a] | Pre-rRNA processing protein UTP22 [a] |
| exon | 13 | 337 | - | NANOZOOG6896 | | | |
| exon | 398 | 429 | - | NANOZOOG6896 | | | |
| gene | 516 | 3036 | + | NANOZOOG6897 | | | |
| transcript | 516 | 3036 | + | NANOZOOG6897 | NANOZOOT6897.1 | *UTP22* | U3 small nucleolar RNA-associated protein 2[illegible] |
| exon | 516 | 547 | + | NANOZOOG6897 | | | |
| exon | 606 | 727 | + | NANOZOOG6897 | | | |
| exon | 782 | 2414 | + | NANOZOOG6897 | | | |
| exon | 2471 | 2692 | + | NANOZOOG6897 | | | |
| exon | 2758 | 3036 | + | NANOZOOG6897 | | | |
| transcript | 538 | 2370 | + | NANOZOOG6897 | NANOZOOT6897.2 | *UTP22* | U3 small nucleolar RNA-associated protein 2[illegible] |
| exon | 538 | 727 | + | NANOZOOG6897 | | | |
| exon | 782 | 2370 | + | NANOZOOG6897 | | | |
| gene | 4438 | 8540 | - | NANOZOOG6898 | | | |
| transcript | 4438 | 8540 | - | NANOZOOG6898 | NANOZOOT6898.1 | *H2A* | Histone H2A |
| exon | 4438 | 7931 | - | NANOZOOG6898 | | | |
| exon | 7997 | 8097 | - | NANOZOOG6898 | | | |
| exon | 8164 | 8230 | - | NANOZOOG6898 | | | |
| exon | 8303 | 8540 | - | NANOZOOG6898 | | | |
| gene | 8726 | 9573 | + | NANOZOOG6899 | | | |
| transcript | 8726 | 9573 | + | NANOZOOG6899 | NANOZOOT6899.1 | *H2B* | Histone H2B |
| exon | 8726 | 8811 | + | NANOZOOG6899 | | | |
| exon | 8893 | 9032 | + | NANOZOOG6899 | | | |
| exon | 9095 | 9204 | + | NANOZOOG6899 | | | |
| exon | 9275 | 9573 | + | NANOZOOG6899 | | | |

[a] Gene code and name for transcript “NANOZOOT6896.1” were identified through Blast search.

**Supplementary Table 4.** Distribution of exon counts in the transcriptome

| **Exons per transcript** | **Transcripts** |
|---|---|
| 1 | 914 |
| 2 | 2237 |
| 3 | 1935 |
| 4 | 1570 |
| 5 | 1283 |
| 6 | 1004 |
| 7 | 759 |
| 8 | 557 |
| 9 | 407 |
| 10 | 288 |
| 11 | 190 |
| 12 | 156 |
| 13 | 109 |
| 14 | 70 |
| 15 | 69 |
| 16 | 38 |
| 17 | 25 |
| 18 | 27 |
| 19 | 17 |
| 20 | 9 |
| 21 | 3 |
| 22 | 5 |
| 23 | 8 |
| 24 | 2 |
| 25 | 3 |
| 26 | 2 |
| 28 | 1 |
| 31 | 1 |
| 32 | 2 |
| total | 11691 |

**Supplementary Table 5.** Examples of lipid and carotenoid metabolism related genes identified in the genome of *Rhodotorula toruloides* CBS 14.

| Gene | Enzyme | EC | Pathway | Contig/scaffold |
|---|---|---|---|---|
| *CDC19* | Pyruvate kinase 1 | 2.7.1.40 | Glycolysis | contig_35_segment0_pilon_pilon |
| *MAE1* | NADP-dependent malic enzyme | 1.1.1.40 | NADPH regeneration | contig_35_segment0_pilon_pilon |
| *MAE2* | NAD-dependent malic enzyme | 1.1.1.38 | Tricarboxylic acid cycle | contig_19_segment0_pilon_pilon |
| *ACL1* | ATP-citrate synthase | 2.3.3.8 | Lipid synthesis | contig_67_segment0_pilon_pilon |
| *ACC1* | Acetyl-CoA carboxylase | 6.3.4.14 | Fatty acid synthesis | scaffold_3_segment0_pilon_pilon |
| *FAS1* | β- subunit of fatty acid synthase complex | 2.3.1.86 | Fatty acid synthesis | scaffold_3_segment0_pilon_pilon |
| *FAS2* | $\alpha$- subunit of fatty acid synthase complex | 2.3.1.86 | Fatty acid synthesis | contig_5_segment0_pilon_pilon |
| *FAS21* | $\alpha$- subunit of fatty acid synthase complex | 2.3.1.86 | Fatty acid synthesis | contig_5_segment0_pilon_pilon |
| *OLE1* | Acyl-CoA desaturase 1 | 1.14.19.1 | Unsaturated fatty acid synthesis | contig_67_segment0_pilon_pilon |
| *ACAD10* | Acyl-CoA dehydrogenase family member 11 | 1.3.99 | Fatty acid beta-oxidation | contig_30_segment0_pilon_pilon |
| *ACAD12* | Acyl-CoA dehydrogenase family member 10 | 1.3.99 | Fatty acid beta-oxidation | contig_36_segment0_pilon_pilon |
| *crtYB* | Bifunctional lycopene cyclase/phytoene synthase | 5.5.1.19 & 2.5.1.32 | Carotenoid synthesis | contig_30_segment0_pilon_pilon |
| *crtI* | Phytoene desaturase | 1.3.99.30 | Carotenoid synthesis | contig_30_segment0_pilon_pilon |
| *BTS1* | Geranylgeranyl pyrophosphate synthase | 2.5.1.- | Carotenoid synthesis | scaffold_3_segment0_pilon_pilon |

**Supplementary Table 6.** Genes that code for the $\alpha$- subunit of the fatty acid synthase complex and their transcripts.

| contig_5_segment0_pilon_pilon | from | to | Sense | Gene ID | Transcript ID | Gene code |
|---|---|---|---|---|---|---|
| gene | 152253 | 164206 | - | NANOZOOG6383 | | |
| transcript | 152253 | 162314 | - | NANOZOOG6383 | NANOZOOT6383.1 | *FAS2* |
| exon | 152253 | 152925 | - | NANOZOOG6383 | | |
| exon | 152980 | 153199 | - | NANOZOOG6383 | | |
| exon | 153256 | 153357 | - | NANOZOOG6383 | | |
| exon | 153415 | 153542 | - | NANOZOOG6383 | | |
| exon | 153599 | 154382 | - | NANOZOOG6383 | | |
| exon | 154441 | 154708 | - | NANOZOOG6383 | | |
| exon | 154771 | 155013 | - | NANOZOOG6383 | | |
| exon | 155073 | 156241 | - | NANOZOOG6383 | | |
| exon | 156307 | 156779 | - | NANOZOOG6383 | | |
| exon | 156840 | 156985 | - | NANOZOOG6383 | | |
| exon | 157046 | 158889 | - | NANOZOOG6383 | | |
| exon | 158951 | 159562 | - | NANOZOOG6383 | | |
| exon | 159625 | 161438 | - | NANOZOOG6383 | | |
| exon | 161499 | 161714 | - | NANOZOOG6383 | | |
| exon | 161777 | 162048 | - | NANOZOOG6383 | | |
| exon | 162228 | 162314 | - | NANOZOOG6383 | | |
| transcript | 153614 | 164206 | - | NANOZOOG6383 | NANOZOOT6383.2 | *FAS2* |
| exon | 153614 | 154382 | - | NANOZOOG6383 | | |
| exon | 154441 | 154708 | - | NANOZOOG6383 | | |
| exon | 154771 | 155013 | - | NANOZOOG6383 | | |
| exon | 155073 | 156241 | - | NANOZOOG6383 | | |
| exon | 156307 | 156779 | - | NANOZOOG6383 | | |
| exon | 156840 | 156985 | - | NANOZOOG6383 | | |
| exon | 157046 | 158889 | - | NANOZOOG6383 | | |
| exon | 158951 | 159562 | - | NANOZOOG6383 | | |
| exon | 159625 | 161438 | - | NANOZOOG6383 | | |
| exon | 161499 | 161714 | - | NANOZOOG6383 | | |
| exon | 161777 | 162048 | - | NANOZOOG6383 | | |
| exon | 164120 | 164206 | - | NANOZOOG6383 | | |
| transcript | 159625 | 161636 | - | NANOZOOG6383 | NANOZOOT6383.3 | *FAS2* |
| exon | 159625 | 159953 | - | NANOZOOG6383 | | |
| exon | 160098 | 161438 | - | NANOZOOG6383 | | |
| exon | 161499 | 161636 | - | NANOZOOG6383 | | |
| gene | 157050 | 159552 | + | NANOZOOG6384 | | |
| transcript | 157050 | 159552 | + | NANOZOOG6384 | NANOZOOT6384.1 | *FAS21* |
| exon | 157050 | 158090 | + | NANOZOOG6384 | | |
| exon | 158965 | 159552 | + | NANOZOOG6384 | | |

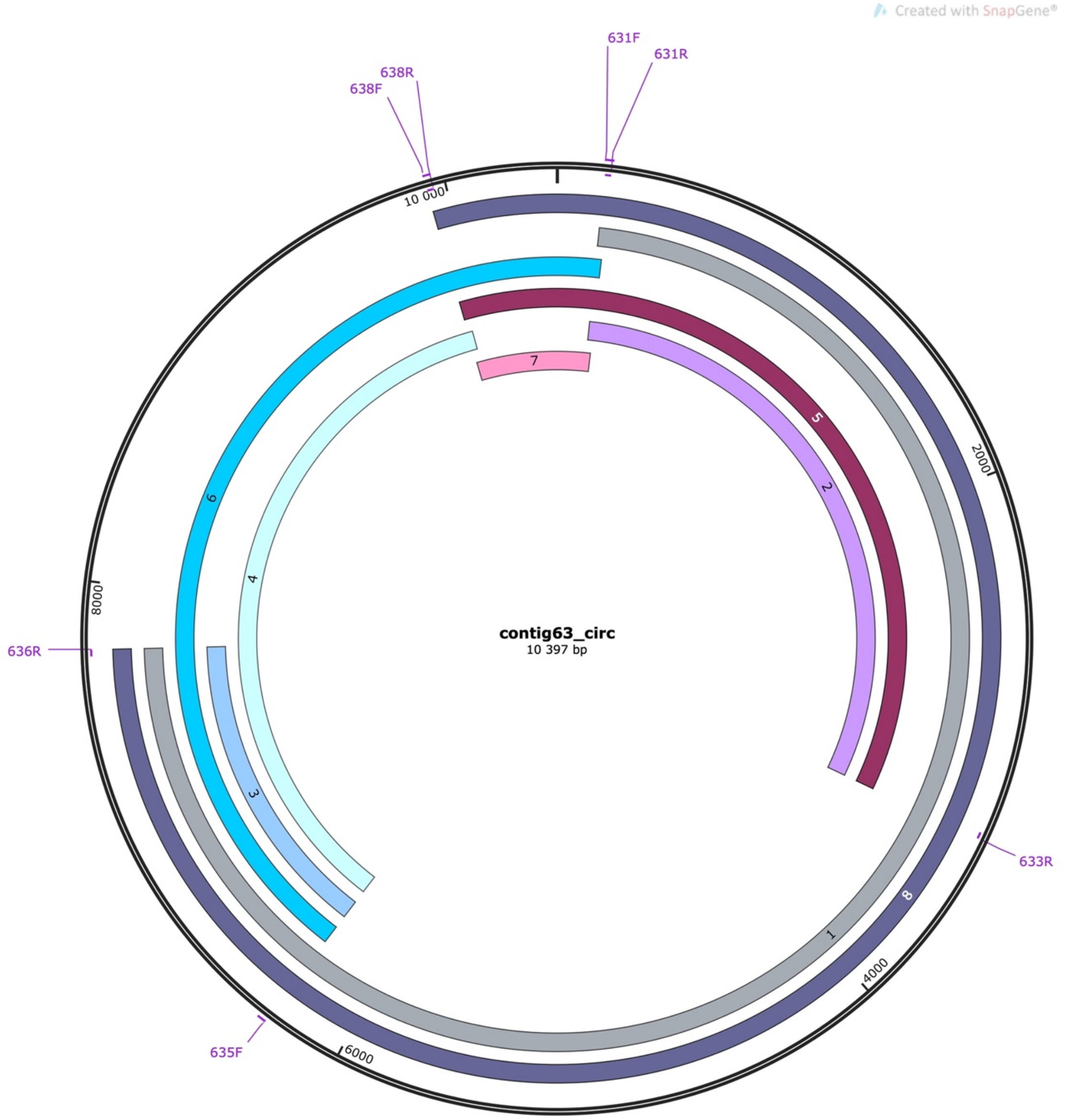


a.

| Fragment | Primer pairs | Expected size (bp) |
|---|---|---|
| 1 | 631F-636R | 7.6 |
| 2 | 631F-633R | 3.2 |
| 3 | 635F-636R | 1.5 |
| 4 | 635F-638R | 3.7 |
| 5 | 638F-633R | 4.9 |
| 6 | 635F-631R | 5.5 |
| 7 | 638F-631R | 1.8 |
| 8 | 638F-636R | 9.4 |

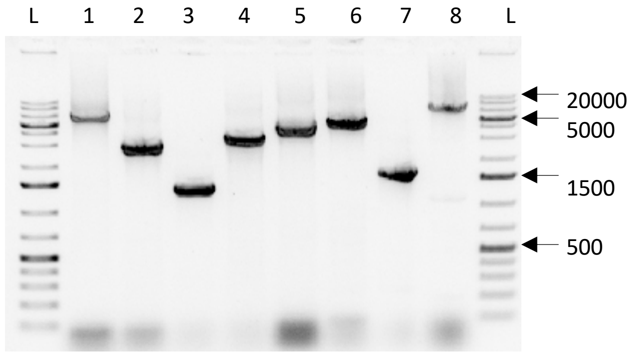


b.

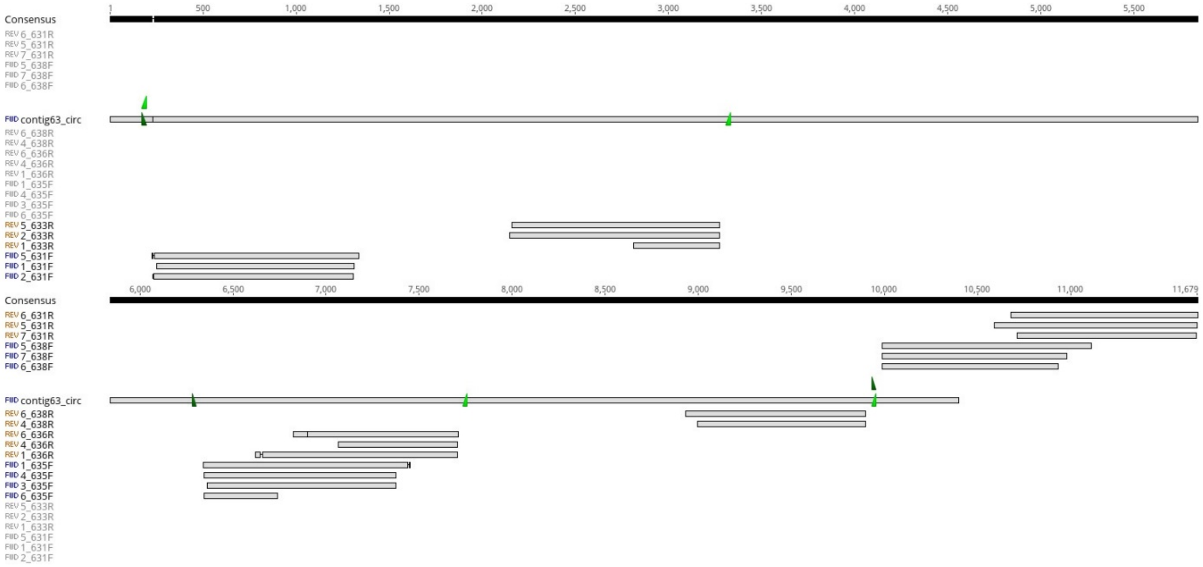


c.

**Supplementary Figure 1.** Validation of the circular structure of Contig 63. (a) Schematic representation of the circular structure of Contig 63, position of primer binding sites and theoretical fragments depending on primer combination. (b) Expected amplicon size (table) and obtained amplicons analyzed by agarose gel electrophoresis using 1% agarose gel. Lane L, 1 Kb plus DNA ladder. (c) Sequence assembly of Contig 63 and amplicon sequences obtained by Sanger sequences. The primer sites within Contig 63 are represented as dark green for forward primers and as light green for reverse primers. The primer name is indicated in the sequence name. The consensus sequence of the contig is represented as a black bar. In the bar representation of each sequence, an average pair-wise identity of 100% is shown in gray and gaps or mismatches as a horizontal line.

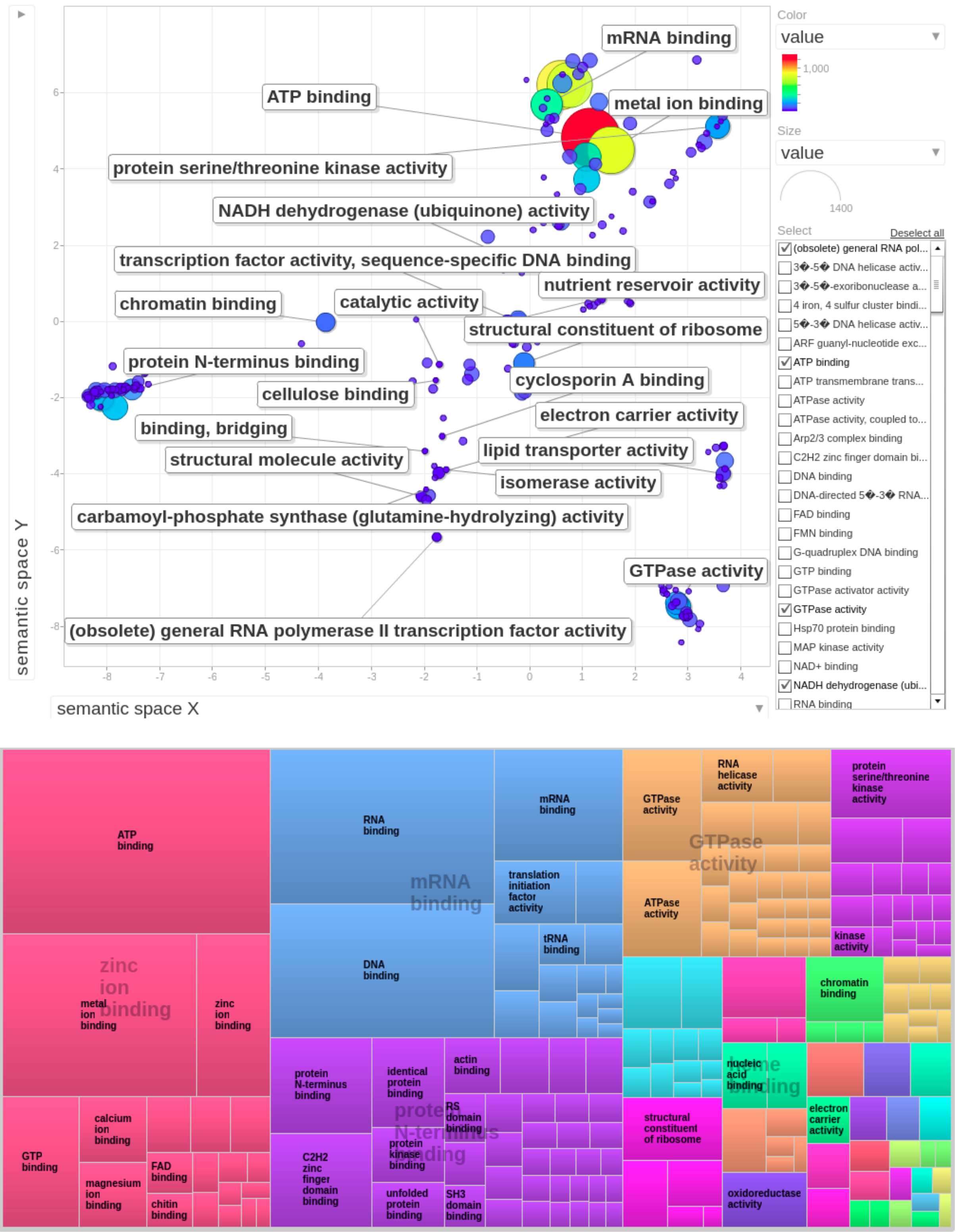


**Supplementary Figure 2.** Gene Ontology (GO) term summary related to the GO topic: molecular functions

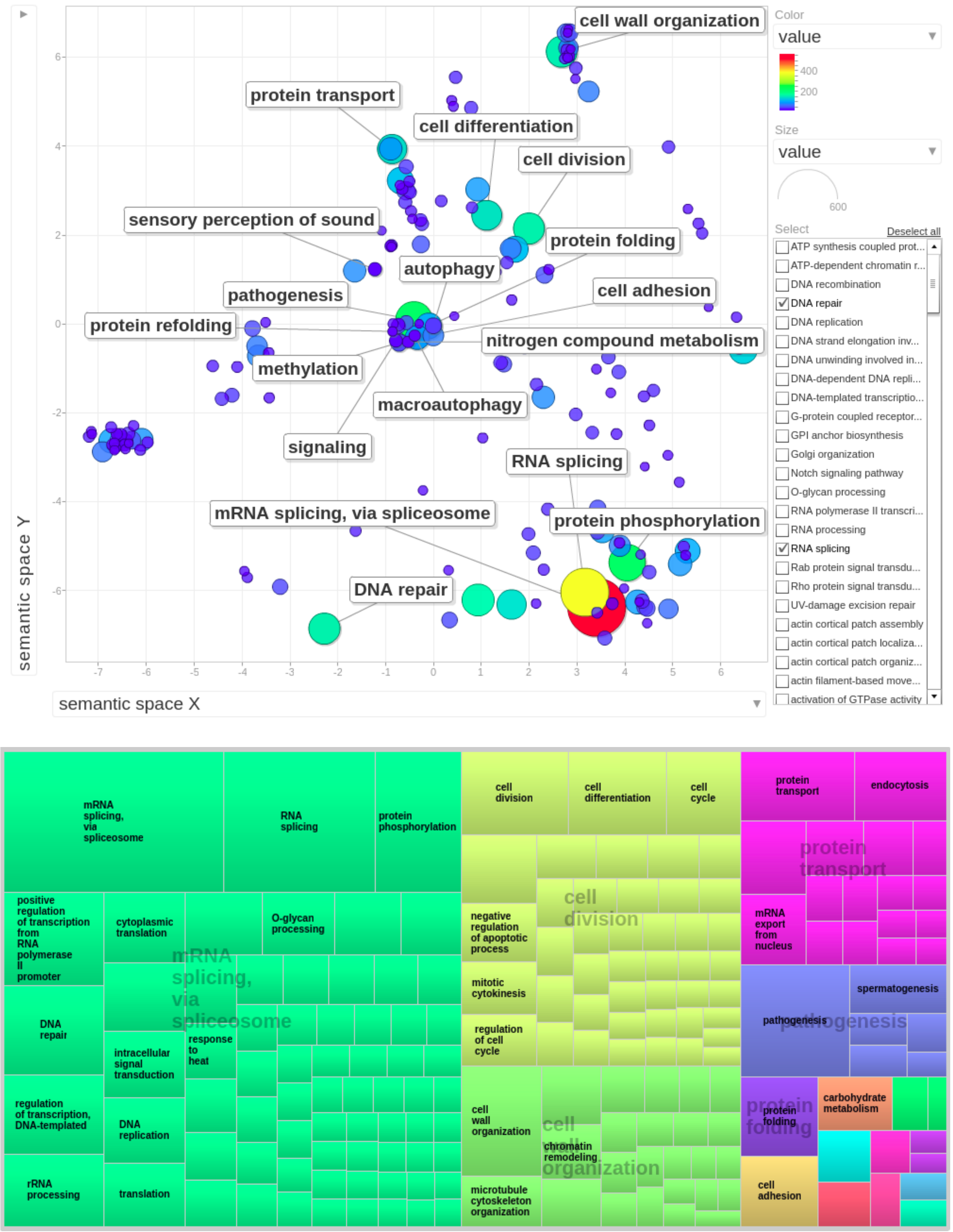


**Supplementary Figure 3.** Gene Ontology (GO) term summaries belonging to the GO topic: biological processes

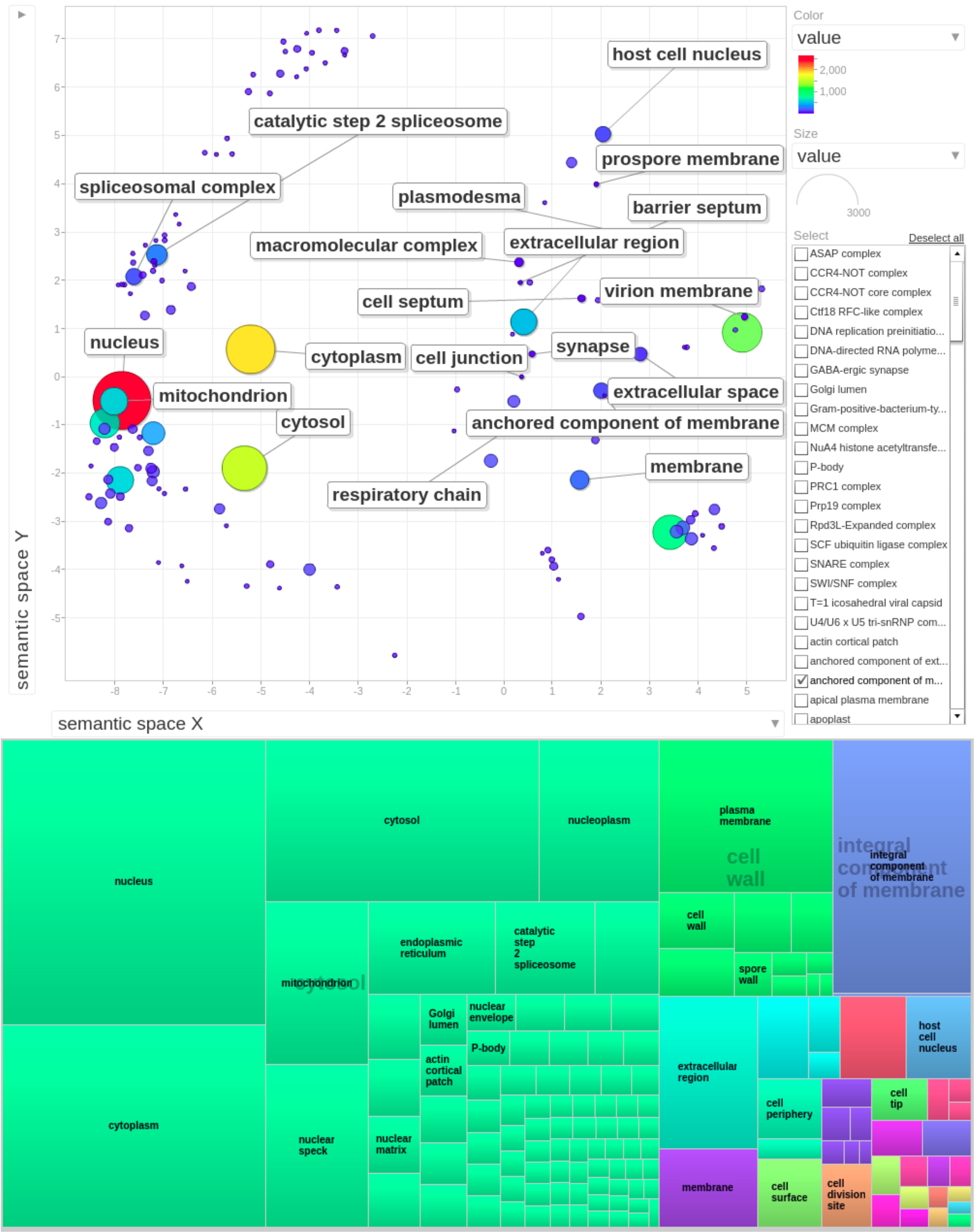


**Supplementary Figure 4.** Gene Ontology (GO) term summaries belonging to the GO topic: cellular components

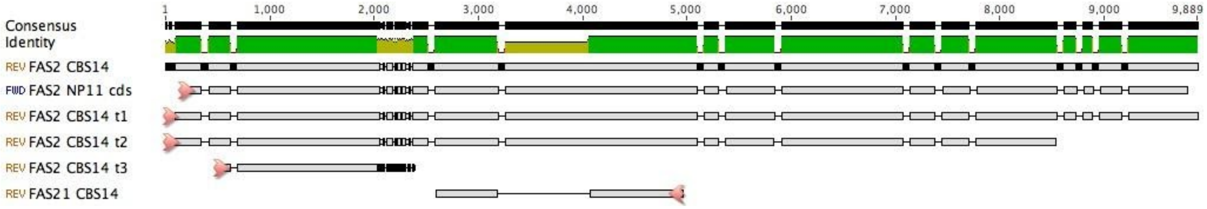


**Supplementary Figure 5. Sequence** alignment of *FAS2* and *FAS21* from *Rhodotorula toruloides* CBS 14 and their transcripts and *FAS2* from *Rhodotorula toruloides* NP11. *FAS2*-genes encode for the $\alpha$- subunit of the fatty acid synthase complex. The alignment was built using the sequences of the three *FAS2* transcripts obtained using StringTie software and guided by the MetaEuk annotation, and *FAS2* from *Rhodotorula toruloides* NP11. The *FAS2* gene was included to identify introns. The consensus sequence is represented as a black bar. The identity bar represents pairs as green for 100% mean pairwise identity, green-brown for at least 30% and under 100%, and red for below 30%. In the bar representation of each sequence, pairs with 100% mean pairwise identity are gray and the gaps are represented as a horizontal line. On the opposite strand, another gene, *FAS21*, was identified by gene annotation using MetaEuk. Transcription direction is indicated with a pink arrow.

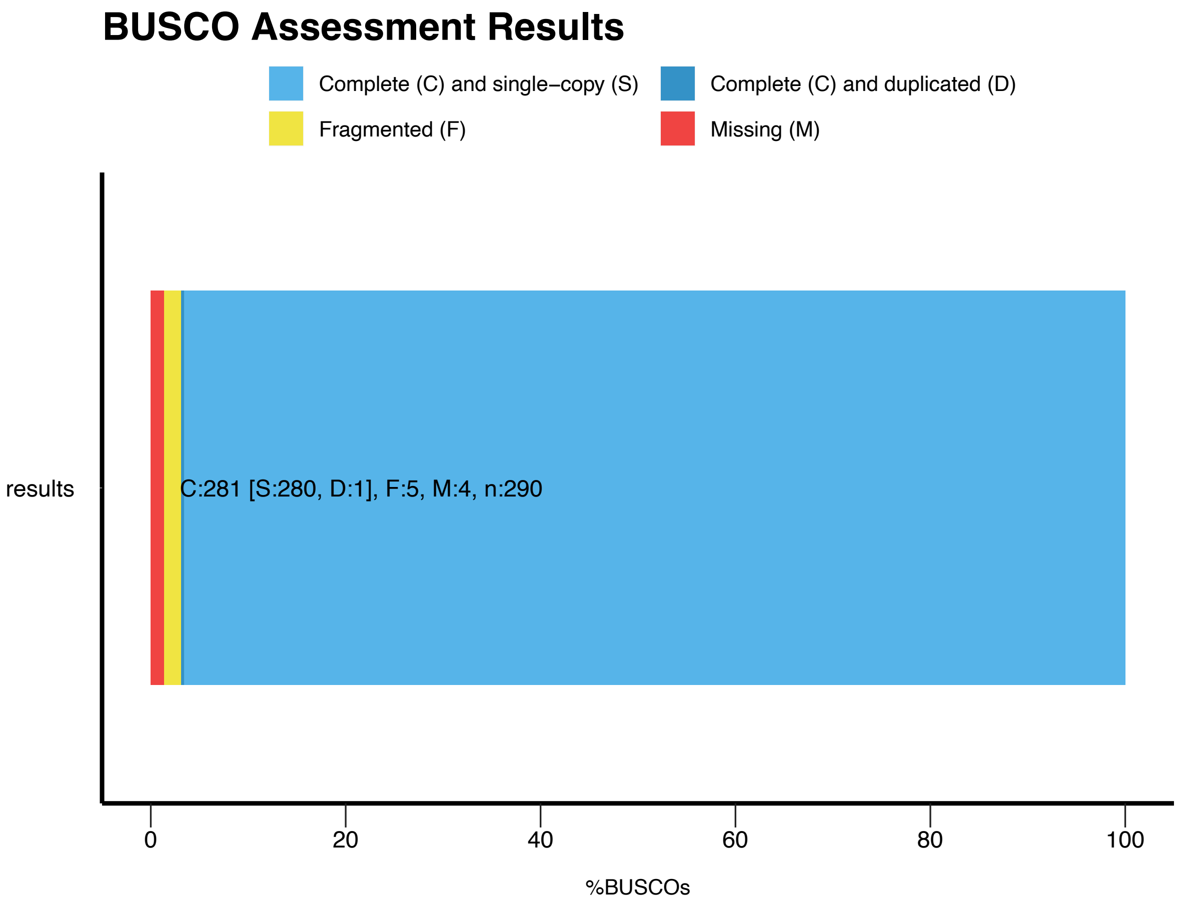


**a.**

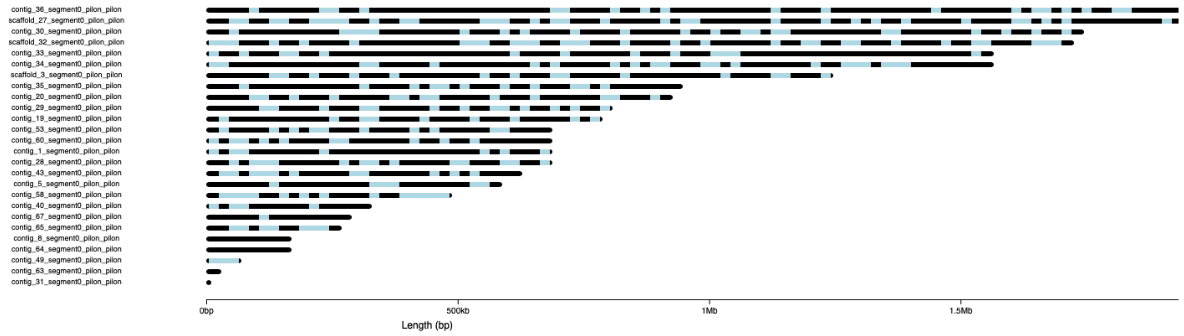


**b.**

**Supplementary Figure 6. Quantitative assessment of the hybrid genome assembly and annotation completeness using Benchmarking Universal Single-Copy Orthologs (BUSCO).** (a) Completeness of the hybrid genome assembly. (b) Graphical overview of the identified near-universal single-copy orthologs.
